# The Stickland fermentation precursor trans-4-hydroxyproline differentially impacts the metabolism of *Clostridioides difficile* and commensal *Clostridia*

**DOI:** 10.1101/2021.11.12.468462

**Authors:** A.D. Reed, J.R. Fletcher, Y.Y. Huang, R. Thanissery, A.J. Rivera, R.J. Parsons, A. Stewart, D.J. Kountz, A. Shen, E.P. Balskus, C.M. Theriot

**Affiliations:** Department of Population Health and Pathobiology, College of Veterinary Medicine, North Carolina State University, Raleigh, NC USA; Molecular Education, Technology and Research Innovation Center, North Carolina State University; Department of Chemistry and Chemical Biology, Harvard University, Cambridge, MA USA; Department of Molecular Biology and Microbiology, Tufts University School of Medicine, Boston, Massachusetts, USA

**Keywords:** *Clostridioides difficile*, hydroxyproline, amino acids, *Clostridia*, Stickland fermentation, colonization resistance

## Abstract

An intact gut microbiota confers colonization resistance against *Clostridioides difficile* through a variety of mechanisms, likely including competition for nutrients. Recently, proline was identified as an important environmental amino acid that *C. difficile* uses to support growth and cause significant disease. A post-translationally modified form, trans-4-hydroxyproline, is highly abundant in collagen, which is degraded by host proteases in response to *C. difficile* toxin activity. The ability to dehydrate trans-4-hydroxyproline via the HypD glycyl radical enzyme is wide-spread amongst gut microbiota, including *C. difficile* and members of the commensal *Clostridia*, suggesting that this amino acid is an important nutrient in the host environment. Therefore, we constructed a *C. difficile* Δ*hypD* mutant and found that it was modestly impaired in fitness in a mouse model of infection, and was associated with an altered microbiota when compared to mice challenged with the wild type strain. Changes in the microbiota between the two groups were largely driven by members of the *Lachnospiraceae* family and the *Clostridium* genus. We found that *C. difficile* and type strains of three commensal *Clostridia* had significant alterations to their metabolic gene expression in the presence of trans-4-hydroxyproline *in vitro*. The proline reductase (*prd*) genes were elevated in *C. difficile*, consistent with the hypothesis that trans-4-hydroxyproline is used by *C. difficile* to supply proline for fermentation. Similar transcripts were also elevated in some commensal *Clostridia* tested, although each strain responded differently. This suggests that the uptake and utilization of other nutrients by the commensal *Clostridia* may be affected by trans-4-hydroxyproline metabolism, highlighting how a common nutrient may be a signal to each organism to adapt to a unique niche. Further elucidation of the differences between them in the presence of hydroxyproline and other key nutrients will be important to determining their role in nutrient competition against *C. difficile.*

## Introduction

*Clostridioides difficile* infection (CDI) is the cause of significant morbidity and mortality and is responsible for over 4.8 billion dollars in excess medical costs each year[1,2]. The current front-line treatment for CDI is the antibiotic vancomycin, which can resolve CDI[3]. However, 20-30% of patients will experience a recurrence of CDI within 30 days, and 40-60% of the patients who have experienced one recurrence will have multiple recurrences[4,5]. The use of antibiotics, including vancomycin, is a major risk factor for CDI due to their effect on the gut microbiota, which causes a loss of colonization resistance against *C. difficile*[6–8]. Colonization resistance, or the ability of the gut microbiota to defend against colonization by gastrointestinal pathogens such as *C. difficile* has many potential mechanisms, including the production of inhibitory metabolites and competition for nutrient sources[9–11]. Conversely, *C. difficile* toxin activity is associated with altered recovery of the gut microbiota, as well as liberation of numerous sugars and peptides/amino acids *in vivo*[12–15]. However, it is unknown if *C. difficile* has a hierarchy of preferred nutrient sources in a host, or whether members of the microbiota also utilize similar nutrients, and if they do, whether their use contributes to colonization resistance.

Much of the research on colonization resistance against *C. difficile* has focused on the effects of secondary bile acids produced by the gut microbiota[16–20]. While secondary bile acid metabolism is an important contributor, other factors such as competition for nutrients are also likely to play a role. For example, colonization of a host by non-toxigenic *C. difficile* can prevent colonization by toxigenic *C. difficile*, indicating that bacteria with similar nutritional needs can occupy an exclusive niche[21,22]. In addition, the increased amount of succinate available in the antibiotic treated gut promotes expansion by *C. difficile*, indicating that the depletion of the microbiota that occurs after antibiotic use creates a beneficial environment for *C. difficile* colonization and expansion[23]. Metabolic and transcriptomic analysis have also shown that the availability of amino acids and other nutrients is very important in the early stage of CDI[15,23,24]. Additionally, the degradation of collagen by host proteases that is induced by *C. difficile* toxin activity may be a source of peptides and amino acids to support *C. difficile* growth through the course of infection[14,15].

*C. difficile* uses proline as an electron acceptor for Stickland fermentation for energy production and regeneration of NAD^+^, yet it does not grow well without the presence of proline and other amino acids important to Stickland fermentation, therefore it must compete for them within the host environment[25–27]. The concentration of proline in media affects expression of genes in the *prd* operon, which encodes proline reductase and accessory proteins, with maximal expression observed when proline content is high[26]. In addition, the availability of proline and branched chain amino acids in the gut correlates with increased susceptibility to *C. difficile* in a mouse model of infection[13]. When a *C. difficile prdB* mutant that was unable to utilize proline as an energy source was tested in a mouse model of CDI, it was less fit *in vivo* and resulted in less toxin (TcdB) in stools when compared to mice challenged with wild type *C difficile*[13]. In addition, the presence of some commensal *Clostridia* causes an increase in the reliance of *C. difficile* on proline fermentation [28]. This indicates that *C. difficile* may compete with some commensal *Clostridia* for proline in the gut. *C. difficile* also has a competitive advantage over the commensals *Clostridium scindens, Clostridium hylemonae*, and *Clostridium hiranonis* in a rich medium, although the extent to which this is due to the ability of *C. difficile* to ferment proline is unknown[16].

*Trans*-4-hydroxy-L-proline (hydroxyproline or hyp) is a derivative of proline that has been post-translationally modified by the host via prolyl-4-hydroxylase, and is a significant component of the highly abundant host protein collagen. Recently, we have shown that inflammation resulting from *C. difficile* toxin activity leads to increased expression of host matrix metalloproteinases and subsequent degradation of collagen, likely supplying *C. difficile* with hydroxyproline and other Stickland substrates[14]. *C. difficile* can reduce hydroxyproline to proline in a two-step process that requires the glycyl radical enzyme 4-hydroxyproline dehydratase (HypD) and a pyrroline-5-carboxylate reductase (P5CR) encoded by the gene *proC* [28–30]. Homologs of HypD are widespread in the gut microbiome, and a subset of organisms, largely *Clostridia*, that carry the *hypD* gene also encode an adjacent P5CR homolog, indicating that the ability of bacteria to reduce hydroxyproline may be useful in the gut[29]. The widespread presence of HypD and the competitive fitness advantage gained by proline fermentation indicates that the ability to ferment proline may play a significant role in *C. difficile* colonization in the gut[29,31].

In this study, we hypothesized that use of hydroxyproline by *C. difficile* contributes to its fitness *in vivo*. We tested this by examining disease kinetics of wild type (WT) *C. difficile* and a Δ*hypD* mutant in a mouse model of CDI. Mice challenged with the Δ*hypD* mutant had reduced weight loss, less toxin activity, and increased relative abundances of cecal *Lachnospiraceae*, a family which includes many commensal *Clostridia*, as well as members of the *Clostridium* genus. We also show that hydroxyproline affects the transcriptomes of *C. difficile* and three commensal *Clostridia* species (*C. scindens*, *C. hylemonae*, and *C. hiranonis*), though each had unique gene expression profiles, with alterations to pathways for carbohydrate and amino acid utilization among them. Together, these data show that *C. difficile* relies on hydroxyproline metabolism *in vivo* for robust sporulation and toxin production. Further, it identifies numerous metabolic pathways in *C. difficile* and commensal *Clostridia* that are affected by hydroxyproline, and the unique response of each organism indicates that hydroxyproline may act as a nutrient source and a signal to prime them for metabolism of other specific nutrients.

## Methods

### Animals and housing

C57BL/6J WT mice (5–8 weeks old; n = 18 male and n = 18 female) were purchased from Jackson Labs. The food, bedding, and water were autoclaved, and all cage changes were performed in a laminar flow hood. The mice were subjected to a 12 h light and 12 h dark cycle. Mice were housed in a room with a temperature of 70 °F and 35% humidity. Animal experiments were conducted in the Laboratory Animal Facilities located on the NCSU CVM campus. Animal studies were approved by NC State’s Institutional Animal Care and Use Committee (IACUC). The animal facilities are equipped with a full-time animal care staff coordinated by the Laboratory Animal Resources (LAR) division at NCSU. The NCSU CVM is accredited by the Association for the Assessment and Accreditation of Laboratory Animal Care International (AAALAC). Trained animal handlers in the facility fed and assessed the status of animals several times per day. Those assessed as moribund were humanely euthanized by CO2 asphyxiation.

### Mouse model of *C. difficile* infection

The mice were given 0.5 mg/mL cefoperazone in their drinking water for 5 days to make them susceptible to *C. difficile* infection, then plain water for 2 days, after which time they (n = 8, 4 males and 4 females) received 10^5^ spores of either *C. difficile* 630Δ*erm* (WT) or *C. difficile* 630Δ*erm*Δ*hypD* (*ΔhypD*) via oral gavage. One group of mice (n = 8, 4 males and 4 females) received cefoperazone and no *C. difficile* spores (cef) and were used as uninfected controls. Mice were weighed daily and monitored for clinical signs of distress (ruffled fur, hunched posture, slow ambulation, etc.). Fecal pellets were collected 1-, 3-, 5-, and 7-days post challenge and diluted 1:10 w/v in sterile PBS, then serially diluted in 96-well PCR plates and plated onto CCFA for enumeration of vegetative *C. difficile* CFU. The serially diluted samples were then removed from the anaerobic chamber and heated to 65°C for 20 min before being passed back into the chamber. The dilutions were plated onto TCCFA for enumeration of spore CFUs. Additional fecal pellets were collected on days 1-7 and stored at –80°C for later use in toxin activity assays and 16S rRNA sequencing.

At day 7 post challenge, mice were humanely sacrificed, and necropsy was performed. Cecal content was harvested for enumeration of vegetative *C. difficile* and spore CFUs, as well as for toxin activity. Cecal tissue was harvested for 16S rRNA sequencing. Samples for sequencing and toxin activity were immediately flash frozen in liquid nitrogen and stored at −80°C until processing.

Toxin activity in the cecal content was quantified using the Vero Cell cytotoxicity assay[32]. Briefly, the content was diluted 1:10 w/v in sterile PBS, and 10-fold dilutions were added to Vero cells in a 96-well dish for ~16 h. The reciprocal of the lowest dilution in which ~80% of the cells have rounded was reported as the titer.

### Bacterial strain collection and growth conditions

The *C. difficile* strains used in this study were the wild type *C. difficile* 630Δ*erm* (WT) and the mutants *C. difficile* 630Δ*erm*Δ*hypD* (Δ*hypD*), *C. difficile* 630Δ*erm*Δ*p5cr* (Δ*p5cr*), *C. difficile* 630Δ*erm*Δ*hypD∷hypD* (*hypD* complement), and *C. difficile* 630Δ*erm*Δ*p5cr∷p5cr* (*p5cr* complement). All assays using *C. difficile* were started from spore stocks, which were prepared and tested for purity as described previously [32,33]. *C. difficile* spores were maintained on brain heart infusion (BHI) medium supplemented with 100 mg/L L-cysteine and 0.1% taurocholate (T4009, Sigma-Aldrich). Then cultures were started by inoculating a single colony from the plate into BHI liquid medium supplemented with 100 mg/L L-cysteine. The other bacterial strains used in this study were *C. hiranonis* TO 931*, C. hylemonae* TN 271, and *C. scindens* VPI 12708. All strains were maintained on 15% glycerol stocks stored in –80°C until use and were grown in BHI medium supplemented with 100 mg/L L-cysteine. All strains used in this study were grown under 2.5% hydrogen under anaerobic conditions (Coy, USA) at 37°C.

### Growth studies in CDMM

*C. difficile* was grown in a well-established, defined minimal medium (CDMM)[27]. CDMM –pro +hyp had 600 mg/L of *trans*-4-hydroxy-L-proline (Sigma) instead of L-proline. CDMM –pro was used as a negative control. A single colony was inoculated into 5 mL of media and incubated at 37°C for 24 hr, at which point the OD_600_ was measured using a spectrophotometer.

### Construction of *C. difficile* strains

To construct the pMTL-YN1C-*hypD* complementation construct, primer pair YH-P295 and YH-P296 was used to amplify the *hypD* gene (Supplemental Table 1). The resulting PCR product was digested with NotI and XhoI and ligated to pMTLYN1C digested with the same enzymes. The resulting PCR fragments were inserted into pMTLYN1C digested with NotI and XhoI using Gibson assembly [34]. The assembly mixture was transformed into *E. coli* DH5α, and the resulting plasmids were confirmed by sequencing and then transformed into *E. coli* HB101/pRK24.

### Vectors for gene deletion and complementation

To construct the pMTL-YN3-Δ*hypD* allelic exchange construct, vector~1 kb flanking regions of *hypD* (CD630_32820) were PCR amplified using primers YH-P253 and YH-P25 were used to amplify the region 4(upstream of *hypD*,) and primers YH-254 and YH-256 were used to amplify a region downstream of *hypD* using *C. difficile* 630 genomic DNA as the template. The resulting PCR products were used in a PCR splice overlap extension (SOE) reaction with the flanking primers YH-257 and YH259. To construct the pMTL-YN3-Δ*p5cr* allelic exchange construct, primers YH-P258 and YH-P260 were used to amplify a region upstream of *p5cr*, and primers YH-254 and YH-256 were used to amplify a region downstream of *p5cr* in *C. difficile* (Supplemental Table 1). To construct the pMTL-YN3-Δ*p5cr* allelic exchange vector, ~1 kb flanking regions of *p5cr* or *proC* (CD630_32810) were PCR amplified using primers YH-P257 and YH-P258 (upstream) and primers YH-259 and YH-260 (downstream) with *C. difficile* 630 genomic DNA as the template (Supplemental Table 1). All PCRs of flanking regions were carried out using Phusion-HF Master Mix (NEB) according to the manufacturer’s protocol with an annealing temperature (Ta) of 61°C and extension time of 25 sec. gel-purified using illustra GFX PCR DNA and Gel Band Purification Kit (GE Healthcare). 30 ng of each upstream and downstream flanking region of the targeted gene were used as templates for overlap. PCR products were used in a10 μl reactions (Ta of 72°C, extension time of 120s). 7 μl of each overlap PCR mix was used as template in 20 μl extension PCRs (Ta of 61°C, extension time of 60 sec, Phusion-HF master mix) with 0.75 μM of each flanking primer YHP253 and P256 for Δ*hypD*, and YH-P257 and YH-P260 for Δ*proC* to amplify joined flanking regions. The PCR SOE products were gel-purified and digested with AscI and SbfI. The assembly-HF (NEB). was linearized-HF and gel-purified. Each deletion region was ligated into linearized pMTL-YN3 using T4 DNA ligase (NEB) in 10 μl reactions at a 1:9 volume ratio (vector:insert).Ligation reactions were *E. coli* TOP10 cells and plated out onto LB-chloramphenicol (25 μg/mL) agar plates. To construct the pMTL-YN1C-*hypD* complementation construct, the promoter region, ~300bp, upstream of the HypD activase gene (CD630_32830) and *hypD* were separately amplified using primers YH-P229-P230 (Ta of 63°C, extension time of 10 sec) and YH-P231-P232 (Ta of 59°C, extension time of 60 sec), respectively (Supplemental Table 1). To construct the pMTL-YN1C-*proC* complementation construct, primers YH-P233 and YH-P234 was used to amplify the P5CR-encoding gene along with ~200 bp of the upstream region using a Ta of 60°C, extension time of 20 sec (Supplemental Table 1). The resulting PCR products were gel-purified. 3-fold molar excess of each PCR insert was Gibson assembled into 50 ng or 100 ng of StuI-linearized pMTL-YN1C to construct pMTL-YN1C-*hypD* and pMTL-YN1C-*proC*, respectively. Gibson assembly reactions were transformed into *E. coli* DH5α,TOP10 cells and then plated out onto LB-chloramphenicol (25 μg/mL) agar plates. The resulting plasmids were confirmed by Sanger sequencing and then transformed into *E. coli* HB101/pRK24 for conjugation.

### Gene deletions in *C. difficile*

Allele-coupled exchange was used to construct clean deletions of *hypD* and *PC5R* [34]. The recipient *C. difficile* strain 630Δ*erm*Δ*pyrE* (a kind gift from Nigel Minton, c/o Marcin Dembek) was grown for 5-6 hrs in BHIS medium in an anaerobic chamber (Coy, USA) *E. coli* HB101/pRK24 donor strains carrying the appropriate pMTL-YN3 allelic exchange constructs were grown in LB medium containing ampicillin (50 μg/mL) and chloramphenicol (20 μg/mL) at 37°C°, 225 rpm, under aerobic conditions, for 5-6 hrs. Each *E. coli* strain was pelleted at 2,500 rpm for 5 min and transferred into an anaerobic chamber. One milliliter of the *C. difficile* culture was added to each *E. coli* pellet, and 100 μL of the mixture was spotted seven times onto a BHIS plate. The *E. coli* and *C. difficile* mixture was incubated for 13-18 hrs at 37°C anaerobically after which the resulting growth was scraped from the plate into 1 mL phosphate buffered saline (PBS). One hundred microliter aliquots of each suspension were spread onto five BHIS plates containing 10 μg/mL thiamphenicol, 50 μg/mL kanamycin, and 8 μg/mL cefoxitin. The plates were incubated for 3-4 days at 37°C, and transconjugants were passaged onto BHIS plates containing 15 μg/mL thiamphenicol, 50 μg/mL kanamycin, 8 μg/mL cefoxitin, and 5 μg/mL uracil. After selecting for the fastest growing colonies over 2-3 passages, single colonies were re-struck onto CDMM plates, a defined minimal medium, containing 2 mg/mL 5-fluoroorotic acid (FOA) and 5 μg/mL uracil. FOA-resistant colonies that arose were patched onto CDMM plates containing 5-FOA and uracil, and colony PCR was performed to identify clones harboring the desired deletions[35]. (Supplementary Table 1) All 630Δ*erm*Δ*pyrE* mutant strains were complemented with *pyrE* in the *pyrE* locus as described in the next section.

### Complementation in *C. difficile*

*E. coli* HB101/pRK24 donor strains carrying the appropriate complementation construct were grown in LB containing ampicillin (50 μg/mL) and chloramphenicol (20 μg/mL) at 37°C°, 225 rpm, under aerobic conditions, for 6 hrs. [35,36].For complementation in the *pyrE* locus using pMTL-YN1C constructs, *C. difficile* recipient strains were conjugated with either the empty pMTL-YN1C vector or the appropriate pMTL-YN1C complementation vectors as described previously. Transconjugants were then re-struck onto CDMM and incubated for 2-4 days. Colonies that had restored the *pyrE* locus by virtue of their ability to grow on CDMM were re-struck onto CDMM plates before further characterization. All clones were verified by colony PCR. At least two independent clones from each complementation strain were phenotypically characterized.

### Genomic analysis of *hypD* and *p5cr*

This was performed using the gggenes package (version 0.4.0) in R (version 3.6.3) and Geneious as described previously[16]. Briefly, the *hypD* comparison was constructed by first extracting the positional information for *hypD* and *p5cr* from Geneious [40], then obtaining amino acid identity percentage through BLASTp alignments [41] against coding sequences from the reference strain *C. scindens* ATCC 35704. (NCBI accession no. PRJNA508260) This data was visualized using the publicly available gggenes R package [42].

### RNA extraction

*C. difficile, C. scindens, C. hiranonis* and *C. hylemonae* liquid cultures were started from a single colony and grown in either BHI or BHI + 600 mg/L hydroxyproline media for 14 hr before RNA extraction. Cultures were fixed by adding equal volumes of a 1:1 mixture of EtOH and acetone and stored at –80°C for later RNA extraction. For extraction, the culture was thawed, then centrifuged at 10,000 rpm for 10 min at 4°C. The supernatant was discarded and the cell pellet resuspended in 1 mL of 1:100 BME: (beta-mercaptoethanol)H_2_O, then spun down at 14,000 rpm for 1 min. For RNA to be used for qRT-PCR, the cell pellet was resuspended in 0.3 mL of lysis buffer from the Ambion RNA purification kit (AM1912, Invitrogen) then sonicated while on ice for 10 pulses of 2 sec with a pause of 3 sec between each pulse. Extraction was then performed following the manufacturers protocol from the Ambion RNA purification kit. For RNA to be used for RNA-seq, the cell pellet was resuspended in 1 mL of Trizol (Thermofisher F) and incubated at room temperature for 15 min. 200 μL of chloroform (Sigma-Aldrich) was added, the solution was inverted rapidly for 20 sec, then incubated at room temperature for 15 min and centrifuged at 14,000 rpm for 15 min at 4°C. The aqueous phase was mixed with 96% ethanol and the extraction was performed using the Direct-zol RNA Miniprep Plus following the manufacturer’s instructions, including an on-column DNase I treatment (R2071, Zymo Research).

### Reverse transcription and quantitative real-time PCR

Reverse transcription and quantitative real-time PCR (qRT-PCR) was performed as described previously [15]. Briefly, RNA was depleted by using Turbo DNase according to the manufacturer’s instructions (AM2238, Invitrogen. The DNase-treated RNA was then cleaned using an RNA clean up kit (R1019, Zymo) according to manufacturer’s instructions and DNA depletion was verified by amplifying 1 μL of RNA in a PCR reaction. The DNA depleted RNA was used as the template for reverse transcription performed with Moloney murine leukemia virus (MMLV) reverse transcriptase (M0253, NEB). The cDNA samples were then diluted 1:4 in water and used in quantitative real-time PCR with genespecific primers using SsoAdvanced Universal Sybr green Supermix (1725271, Bio-Rad) according to the manufacturer’s protocol (Supplementary Table 1). Amplifications were performed in technical triplicates, and copy numbers were calculated using a standard curve and normalized to that of a housekeeping gene. *gyrA* was the housekeeping gene used for *C. scindens*, while *rpoC* was used for *C. difficile, C. hiranonis* and *C. hylemonae.*

### RNAseq

Sequencing of RNA derived from *in vitro* cultures was performed at the Roy J. Carver Biotechnology Center at the University of Illinois at Urbana-Champaign. Ribosomal RNA was removed from the samples using the RiboZero Epidemiology Kit. (Illumina). RNAseq libraries were prepped with the TruSeq Stranded mRNA Sample Prep Kit (Illumina), though poly-A enrichment was omitted. Library quantification was done via qPCR, and the samples were sequenced on one lane for 151 cycles from each end of the fragments on a NovaSeq 6000 using a NovaSeq S4 reagent kit. The FASTQ files were generated and demultiplexed using the bcl2fastq v2.20 Conversion Software (Illumina). Raw paired Illumina reads were imported into Geneious 10.2.6, where adapters were removed using BBDuk with a Kmer length of 27. The reads were mapped to the *C. difficile* 630*Δerm* genome (NCBI accession no. NC_009089.1), the *C. hiranonis* DSM 13275 genome (NCBI accession no. GCA_008151785.1), the *C. hylemonae* DSM 15053 genome (NCBI accession no. PRJNA523213), or the *C. scindens* ATCC 35704 genome (NCBI accession no. PRJNA508260) using BBMap with a Kmer length of 10 and no other changes to the default settings. Differential expression analysis between the two conditions was performed using DESeq2 and if genes had an adjusted p-value of <0.05 and ± 1 log_2_ fold change, they were considered differentially expressed. Visualization of differentially enriched genes for each organism was performed using pheatmap (version 1.0.12), ggplots (version 3.3.4), and ggpubr (version 0.4.0.999) within R (version 3.6.3). Some of the differentially enriched genes were hypothetical proteins, those results were removed before the figures were visualized. Full RNAseq data is available in Supplemental Data File 4.

### Metabolomics data analysis. *Mass spectrometry data acquisition*

Samples were diluted 1:100 (10 μL sample, 990 μL water) and transferred to an autosampler vial for analysis by UPLC-MS. For quantification of amino acids, a certified reference material amino acid mix solution (*Trace*-CERT, Sigma) was diluted to achieve a 100 μM working standard solution. Ten calibration standards ranging from 100 μM to 250 nM were prepared by serially diluting the working standard solution. For quantification of 4-hydroxyproline and 5-aminovaleric acid, certified reference material (Sigma) for each was suspended in water to achieve a 1 mg/mL solution which were combined and diluted to achieve a 50 μg/mL working standard solution. Ten calibration standards ranging from 50 μg/mL to 25 ng/mL were prepared by serially diluting the working standard solution. The analysis was performed using a Thermo Vanquish UPLC instrument (Thermo Fisher Scientific, Germering, Germany) coupled to a Thermo Orbitrap Exploris 480 mass spectrometer (Thermo Fisher Scientific, Breman, Germany) with a heated electrospray ionization (HESI) source. Chromatographic separation was achieved on a Waters BEH Amide column (2.1 x 100 mm, 1.8 μM) maintained at 45°C. The following linear gradient of mobile phase A (H_2_O + 0.1% FA) and mobile phase B (MeCN + 0.1% FA) was used: 0-0.1 min (99%B, 0.4 mL/min), 0.1-7 min (99-30%B, 0.4 mL/min), 7-10 min (99%B, 0.4 mL/min). Samples were analyzed (2 μL injections) in positive ion mode (spray voltage 3.5 kV, ion transfer tube temperature 300°C, vaporizer temperature 350°C, sheath gas 50 a.u., aux gas 10 a.u., sweep gas 1 a.u.) with a mass range of m/z 60-1000. MS1 data was collected with a resolving power of 60,000 and an AGC target of 1e6 and ddMS2 data was collected with a resolving power of 30,000, cycle time of 0.6 s, AGC target of 4e5 and stepped HCD collision energy (30, 50, 150). The full data set was acquired in a randomized fashion with water blanks and system suitability samples (QReSS, Cambridge Isotope Laboratories) collected every 10 samples.

### Targeted data processing

Peak integration and amino acid quantification were performed in Skyline^1^. Individual standard curves for each of the 15 amino acids plus hydroxyproline and 5-aminovalerate were constructed using extracted ion chromatogram peak areas from MS1 data and the slope of each curve was calculated using a linear curve fit and a 1/(x *x) weighting. MS2 data was utilized to validate amino acid annotations, particularly to differentiate valine and 5-aminovaleric acid. The concentrations in the study samples were calculated in an identical manner relative to the regression line. Calibration curves for each of the amino acids had R^2^ values ranging from 0.9919 to 0.9994 for the linear range of 0.25 to 100 μM. Calibration curves for 4-hydroxyproline and 5-aminovaleric acid had R^2^ values of 0.9948 to 0.9997, respectively, for the linear range of 0.025 to 50 μg/mL[44].

### 16S rRNA bacterial sequencing

Fecal and cecal samples were sequenced by the University of Michigan Microbial Systems Molecular Biology Laboratory using the Illumina MiSeq platform. Microbial DNA was extracted from the fecal and cecal samples using Mag Attract Power Microbiome kit (Mo Bio Laboratories, Inc.). A dual-indexing sequencing strategy was used to amplify the V4 region of the 16S rRNA gene [65]. Each 20-μlL PCR mixture contained 2 μL of 10X Accuprime PCR buffer II (Life Technologies, CA, 1 USA), 0.15 μL of Accuprime high-fidelity polymerase (Life Technologies, CA, USA), 5 μl of a 4.0 μM primer set, 3 μL DNA, and 11.85 μlL sterile nuclease free water. The template DNA concentration was 1 to 10 ng/μlL for a high bacterial DNA/host DNA ratio. The PCR conditions were as follows: 2 min at 95°C, followed by 30 cycles of 95°C for 20 sec, 55°C for 15 sec, and 72°C for 5 min, followed by 72°C for 10 min.

Libraries were normalized using a Life Technologies SequalPrep normalization plate kit as per manufacturer’s instructions for sequential elution. The concentration of the pooled samples was determined using the Kapa Biosystems library quantification kit for Illumina platforms (Kapa Biosystems, MA, USA). Agilent Bioanalyzer high sensitivity DNA analysis kit (Agilent CA, USA) was used to determine the sizes of the amplicons in the library. The final library consisted of equal molar amounts from each of the plates, normalized to the pooled plate at the lowest concentration. Sequencing was done on the Illumina MiSeq platform, using a MiSeq reagent kit V2 (Ilumina, CA, USA) with 500 cycles according to the manufacturer’s instructions, with modifications [65]. Sequencing libraries were prepared according to Illumina’s protocol for preparing libraries for sequencing on the MiSeq (Ilumina, CA, USA) for 2 or 4 nM libraries. PhiX and genomes were added in 16S amplicon sequencing to add diversity. Sequencing reagents were prepared according to the Schloss SOP (https://www.mothur.org/wiki/MiSeq_SOP#Getting_started), and custom read 1, read 2 and index primers were added to the reagent cartridge. FASTQ files were generated for paired end reads.

### Community microbial sequencing analysis

Analysis of the V4 region of the 16S rRNA gene was performed in the statistical programming environment R using the DADA2 package (version 1.14.1) [45]. Forward/reverse pairs were trimmed and filtered, with forward reads truncated at 240 nt and reverse reads truncated at 200 nt. No ambiguous bases were allowed, and each read was required to have less than two expected errors based on their quality score. Error-corrected amplicon sequence variants (ASVs) were independently inferred for the forward and reverse reads of each sample and then read pairs were merged to obtain final ASVs. Chimeric ASVs were identified and removed. For taxonomic assignments, ASVs were compared to the Silva v132 database (https://zenodo.org/record/1172783). The R package phyloseq (version 1.30) was used to further analyze and visualize data[46]. Inverse Simpson was used to calculate alpha diversity and Kruskal Wallis was used to determine statistical significance between treatment groups. Relative abundance was calculated using phyloseq and visualized using Prism 7.0c, and differential-abundance analysis between the ΔhypD and WT treatment groups was performed using the Aldex2 package (version 1.18.0) and visualized using the ggplots2 package (version 3.3.4) [37–39].

### Statistical analysis

Statistical tests were performed using Prism version 7.0c for Mac OSX (GraphPad Software, La Jolla, CA, USA). Statistical significance was determined using Mann-Whitney for CFUs, spore count, and toxin activity and Kruskal Wallis with Dunns multiple comparisons for mouse weights during infection. Student T Tests with Welch’s Correction was applied to account for multiple comparisons in analyses of other data. Statistical analysis for the 16S and RNAseq results was performed in the R computing environment. Kruskal Wallis was used for alpha diversity, Permanova Adonis in the vegan package (version 2.5-7) was used to test the difference between groups for the beta diversity analysis, and ALDEx2 was used to calculate the differences between treatments using a centered-log-ratio transform of ASV abundance to create an effect size for each ASV[47,49].

## Results

### *C. difficile* requires *hypD* for maximum growth in a defined minimal media supplemented with hydroxyproline

The reduction of hydroxyproline to L-proline is a two-step process requiring the *hypD* and *p5cr* genes (Figure 1A)[30] To test the ability of *C. difficile* to utilize hydroxyproline, wild type, Δ*hypD*, Δ*p5cr* and complemented strains were grown in a defined minimal medium (CDMM) with hydroxyproline substituted for proline at the same concentration (600 mg/L) (Figure 1B). The Δ*hypD* mutant had a significant growth defect in the CDMM –pro +hyp, indicating that HypD is needed to utilize hydroxyproline (p <0.001, Student’s T Test with Welch’s correction). There was no growth defect observed in CDMM –pro +hyp for the Δ*p5cr* mutant, indicating that the particular *proC* homolog tested is not essential for *C. difficile* to utilize hydroxyproline. Interestingly, the WT strain as well as Δ*p5cr* and both complements grew significantly better in CDMM –pro +hyp than they did in CDMM alone (<0.05, Student’s T Test with Welch’s correction). As expected, all strains had very poor growth in CDMM –pro, as proline is essential for *C. difficile* growth.

**Figure 1:**
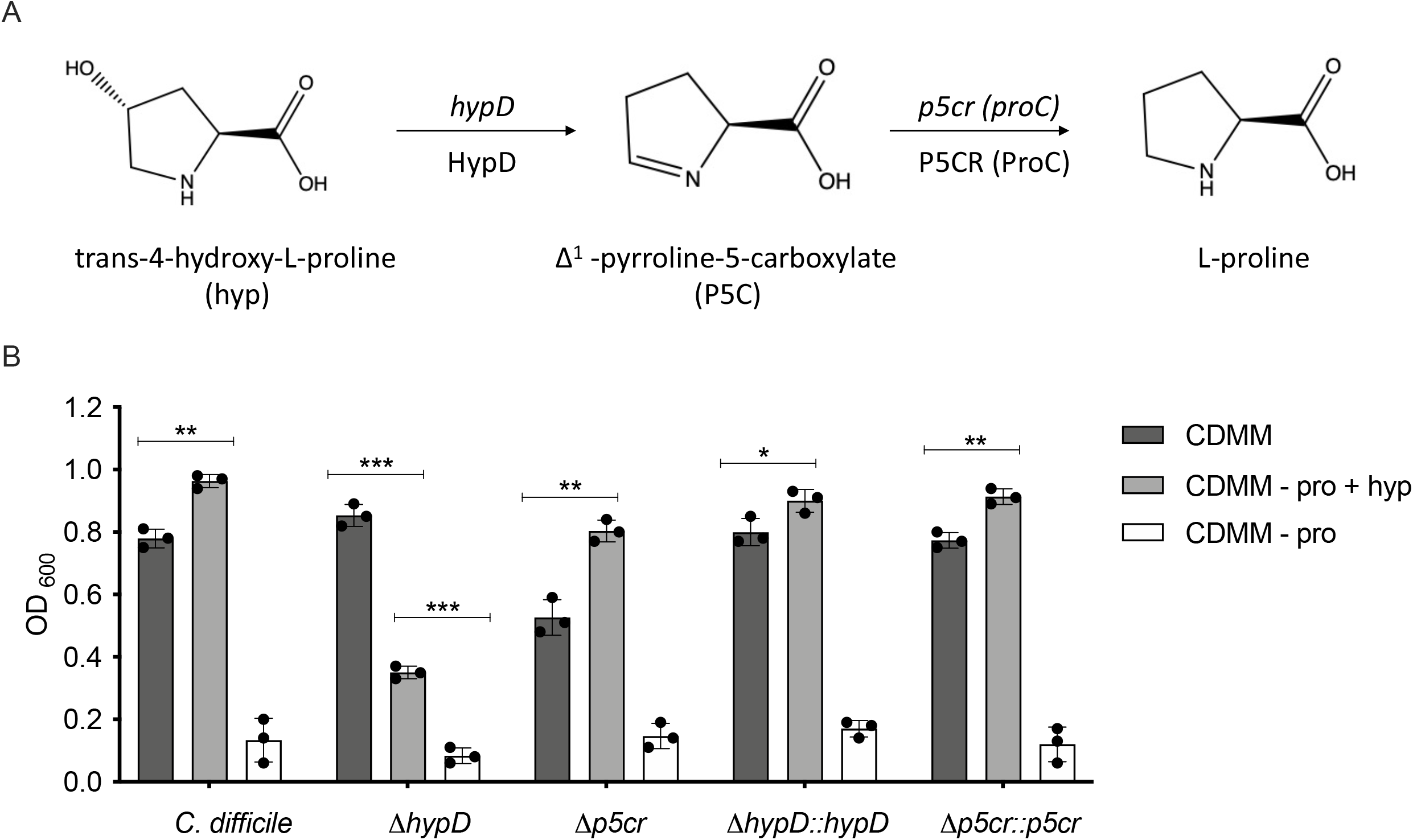
*C. difficile* Δ*hypD* mutant has a growth defect when hydroxyproline is substituted for proline in the growth medium. **A.** Schematic depicting conversion of trans-4-hydroxy-L-proline to L-proline by *hypD* and *p5cr.* **B.** Growth of *C. difficile* 630Δ*erm* WT, the Δ*hypD* and Δ*p5cr* mutants, as well as the complements of both mutants in the defined medium CDMM, in CDMM-proline +hydroxyproline (-pro, +hyp), and CDMM -proline (-pro) Statistical significance was determined using Student’s T Test with Welch’s Correction to account for multiple comparisons (*, P<0.05; **, P<0.01; ***, P<0.001; ****, P<0.0001).

### The presence of *hypD* affects weight loss and toxin activity in a mouse model of CDI

To determine if hydroxyproline utilization is required for colonization and disease, WT C57BL/6J mice (n=8 per group) were challenged with 10^5^ spores of WT *C. difficile* or Δ*hypD*, and colonization and disease progression were measured for 7 days (Figure 2A). There was no significant difference in vegetative *C. difficile* bacterial load in the feces during infection (Figure 2B), but there was a significant decrease in fecal Δ*hypD* spores when compared to the WT group on day 7 post challenge (Figure 2B, p <0.001, Mann-Whitney). On day 7, bacterial enumeration of cecal contents showed no significant difference in the level of *C. difficile* spores (Supplemental Figure 1A and B), while the *C. difficile* vegetative cells were significantly higher in mice challenged with Δ*hypD* (p <0.05, Mann-Whitney). The biggest difference between the groups was seen in the weights of the mice throughout CDI. WT mice weighed significantly less than the cefoperazone control group (cef) on days 3 (p <0.01), 5 (p <0.001), and 7 (p <0.05) post challenge (Figure 2D, Kruskal-Wallis with Dunn’s multiple comparisons). There was no significant difference in weights between the Δ*hypD* group and the cefoperazone control group, indicating that the WT mice had increased clinical signs of disease compared to the Δ*hypD* group. This finding correlated with high toxin activity from the mice in the WT group compared to the Δ*hypD* group on Day 3 post challenge (Figure 2E, p <0.01, Mann-Whitney), although the difference was not significant by Day 7.

**Figure 2:**
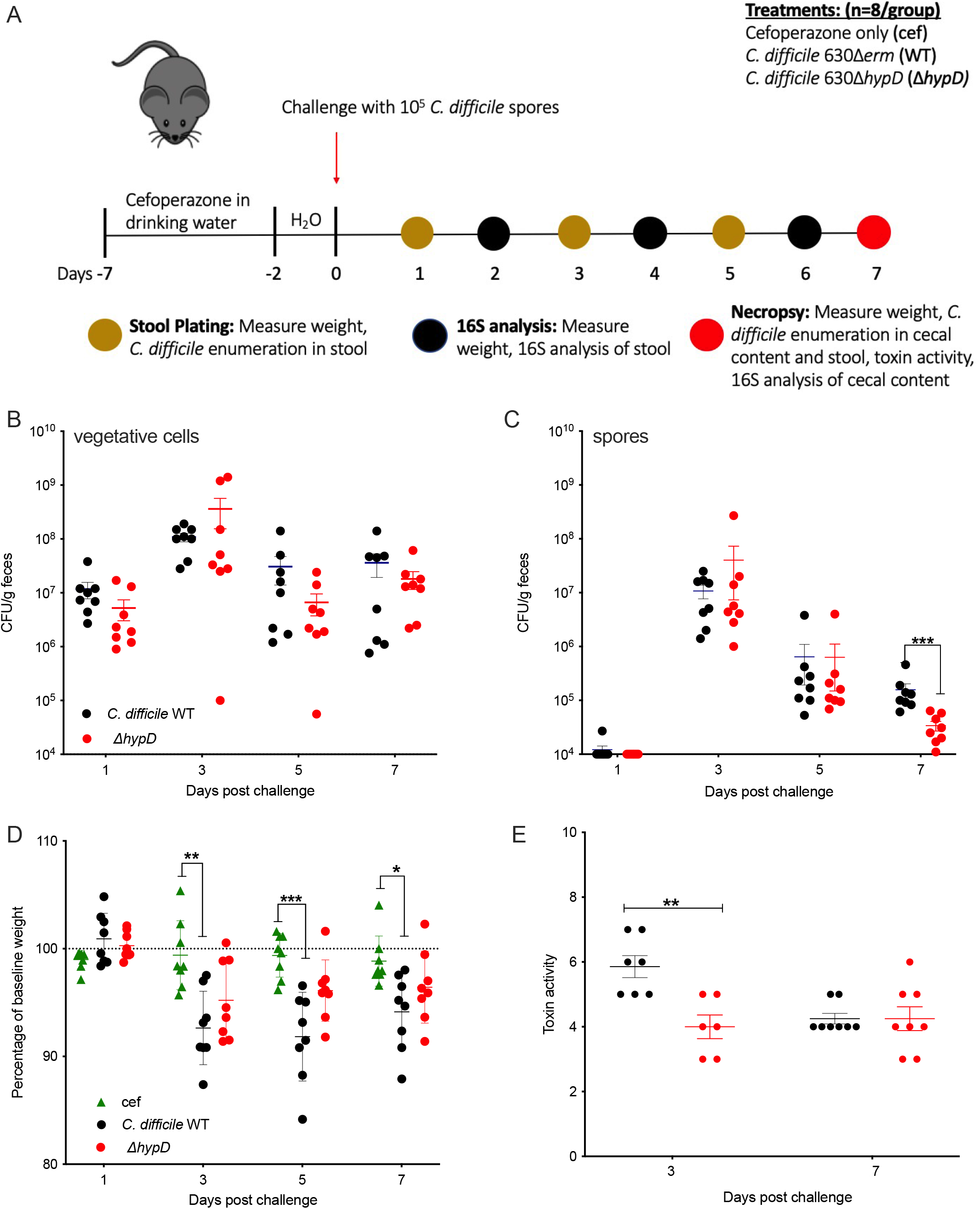
WT *C. difficile* induces more weight loss and toxin activity than the Δ*hypD* mutant in a mouse model of CDI. **A.** Schematic depicting experimental design. All mice (n=24) received the antibiotic cefoperazone in their drinking water. Subsets of mice were orally challenged with *C. difficile* 630Δ*erm* (WT, n=8) or *C. difficile* 630Δ*hypD*Δ*erm* (Δ*hypD*, n=8). The third group of mice were only treated with cefoperazone (cef, n=8). **B-C.** *C. difficile* vegetative cell (**B**) or spore (**C**) CFUs in feces on days 1, 3, 5 and 7 post challenge. **D**. Mouse weights from 1-, 3-, 5- and 7-days post challenge. shown as a percentage of baseline weight for each mouse from day 0 **E**. Toxin activity on 3 and 7 days post challenge. Statistical significance for data shown in **B-E** was determined using Mann-Whitney (*, P<0.05; **, P<0.01; ***, P<0.001; ****, P<0.0001).

### Differences in the microbiota between mice challenged with WT *C. difficile* and Δ*hypD* are driven by members of the *Lachnospiraceae* Family

To elucidate the reason behind the observed differences in CDI between mice challenged with WT and Δ*hypD*, the fecal microbiota of the cef, WT, and Δ*hypD* mice was analyzed through V4 16S rRNA amplicon sequencing on day 0 as well as on days 2, 4 and 6 post challenge (Supplemental Data File 1). The cecal microbiota was analyzed on day 7 post challenge, when necropsy occurred. When the alpha diversity was analyzed in the stool at the Family level, there were significant differences between the cef and Δ*hypD* groups on day 6, and when the cecal microbiota was analyzed on day 7, there were significant differences between all groups (Supplemental Figure 2A, Supplemental Data File 2). When the beta diversity was analyzed using non-metric multi-dimensional scaling analysis (NMDS), there were significant differences between the groups on days 2, 4, 6 and 7 indicating that there was a difference between the three groups after challenge with *C. difficile* (Supplemental Figure 2B). When only the infected groups were analyzed using NMDS, there were significant differences between the WT and Δ*hypD* groups on day 0 and day 7 (Supplemental Figure 2C). On day 0, all three groups had a fecal microbiota dominated by the *Enterococcaceae* (Figure 3A). Day 2 post challenge had the highest relative abundance of *Peptostreptococcaceae*, the family to which *C. difficile* belongs, in both WT and Δ*hypD* mice, which was also when the greatest weight loss was observed (Figure 1D). The amplicon sequence variant (ASV 6) classified as *Peptostreptococcaceae* resolved to *C. difficile*, so this was likely due to the expansion of *C. difficile* in the murine gut microbiota. By day 7 post challenge, the *Lachnospiraceae* family made up a significant percentage of the cecal microbiota for all three groups, with the highest abundance being in the Δ*hypD* group at 73%, while the WT group and the cef group had 60% and 43% abundance of *Lachnospiraceae* respectively (Figure 3A). Differential abundance analysis was calculated between the WT and Δ*hypD* groups using ALDEx2[47]. For each ASV analyzed, ALDEx2 estimates the difference in the centered-log-ratio (a measure of relative abundance) between groups and reports an effect size. The only day that had significant effect sizes for any ASVs was day 7, when the cecal microbiota was analyzed. Although only 3 ASVs were significant, the top 8 ASVs driving differences between the WT and Δ*hypD* microbiotas were examined via NCBI BLAST to determine the identity of each ASV (Figure 3B, Supplemental Data File 1). *Lachnospiraceae bacterium* strain D2 1X 41 and *Clostridium* species MD294 were significantly higher in the WT microbiome than the Δ*hypD* microbiome. *Clostridium* species Clone 49 was significantly higher in the Δ*hypD* microbiota than the WT microbiota. There were also two *Lachnospiraceae* strains, including another ASV that resolved to *Lachnospiraceae bacterium* strain D2 1X 41 as well as *Lachnospiraceae bacterium* DW17, that were higher in the Δ*hypD* microbiome, but did not reach significance. It is unclear why two separate ASVs that resolved to the same strain (*Lachnospiraceae bacterium* strain D2 1X 41) showed such different results in terms of abundance in the WT and Δ*hypD* microbiotas, although it could be due to misassignment, as all 8 ASVs investigated via BLAST matched with multiple strains with high ID, and the highest match was selected to be the identified strain.

**Figure 3:**
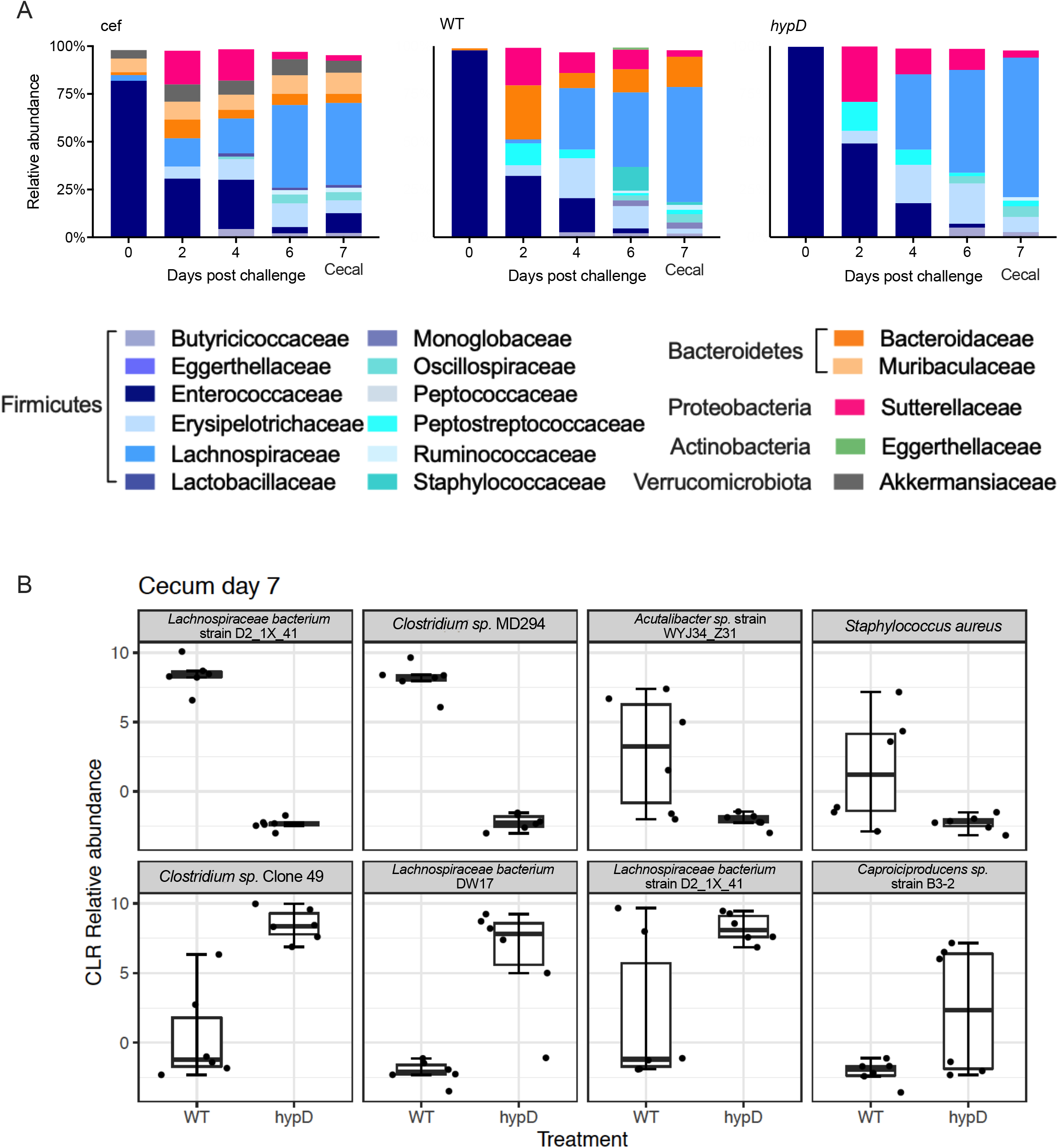
The *Lachnospiraceae* are important when determining the difference between the microbiota of mice challenged with WT and *hypD*. **A.** Relative abundance at the family level for cef, *hypD* and WT fecal microbiome for days 0, 2, 4, and 6 post challenge and the cecal microbiome for day 7 post challenge. **B.** Top 8 ASVs driving differences between *hypD* and WT cecal microbiome on day 7 post challenge. Significant ASVs (q <0.1) are bolded.

Of the top 8 ASVs driving the difference between the WT and Δ*hypD* cecal microbiomes on day 7 post challenge, 5 were either *Clostridium* species or members of the *Lachnospiraceae* family, including all statistically significant ASVs (Figure 3B). This suggests that hydroxyproline may be differentially abundant between the two groups of mice and that members of the *Lachnospiraceae* family and the *Clostridium* genus respond to this by utilizing it for growth. Given the previous work showing that commensal *Clostridia* are important to colonization resistance against *C. difficile*, we next wanted to investigate the response of commensal *Clostridia* to hydroxyproline[16,19].

### Hydroxyproline is utilized by commensal *Clostridia* and *C. difficile* when supplemented into a rich medium

To test for the utilization of hydroxyproline by *C. difficile* and the commensal *Clostridia* strains, WT *C. difficile*, Δ*hypD*, and the commensals *C. hiranonis, C. hylemonae* and *C. scindens* were grown in BHI and in BHI + 600mg/L of hydroxyproline (BHI and BHI +hyp) media for 14 hr, then amino acids and 5-amino-valerate, the product of proline fermentation, were measured using LC/MS. As expected, the BHI +hyp control had significantly higher levels of hydroxyproline than the BHI alone (Figure 4A, p < 0.01, Student’s T Test with Welch’s Correction). WT *C. difficile* utilized hydroxyproline, as did all the commensal *Clostridia*, however, the Δ*hypD* mutant did not (Fig. 4A, p < 0.01, Student’s T Test with Welch’s Correction). There were no significant differences in levels of proline in bacterial cultures grown in BHI compared to BHI +hyp, which is likely explained by the fact that the bacteria were grown in a rich medium and that proline is being metabolized into 5-amino-valerate and other intermediates (Figure 4B). The levels of 5-amino-valerate were higher on average in supernatants for all bacteria grown in BHI or in BHI +hyp than in the media control for either condition, indicating that all strains tested were likely utilizing proline and producing 5-amino-valerate (Figure 4C, Supplemental Data File 3). The levels of 5-amino-valerate were significantly higher for *C. scindens* in BHI +hyp, although it is unclear if the difference is biologically relevant (p < 0.01, Student’s T Test with Welch’s Correction).

**Figure 4:**
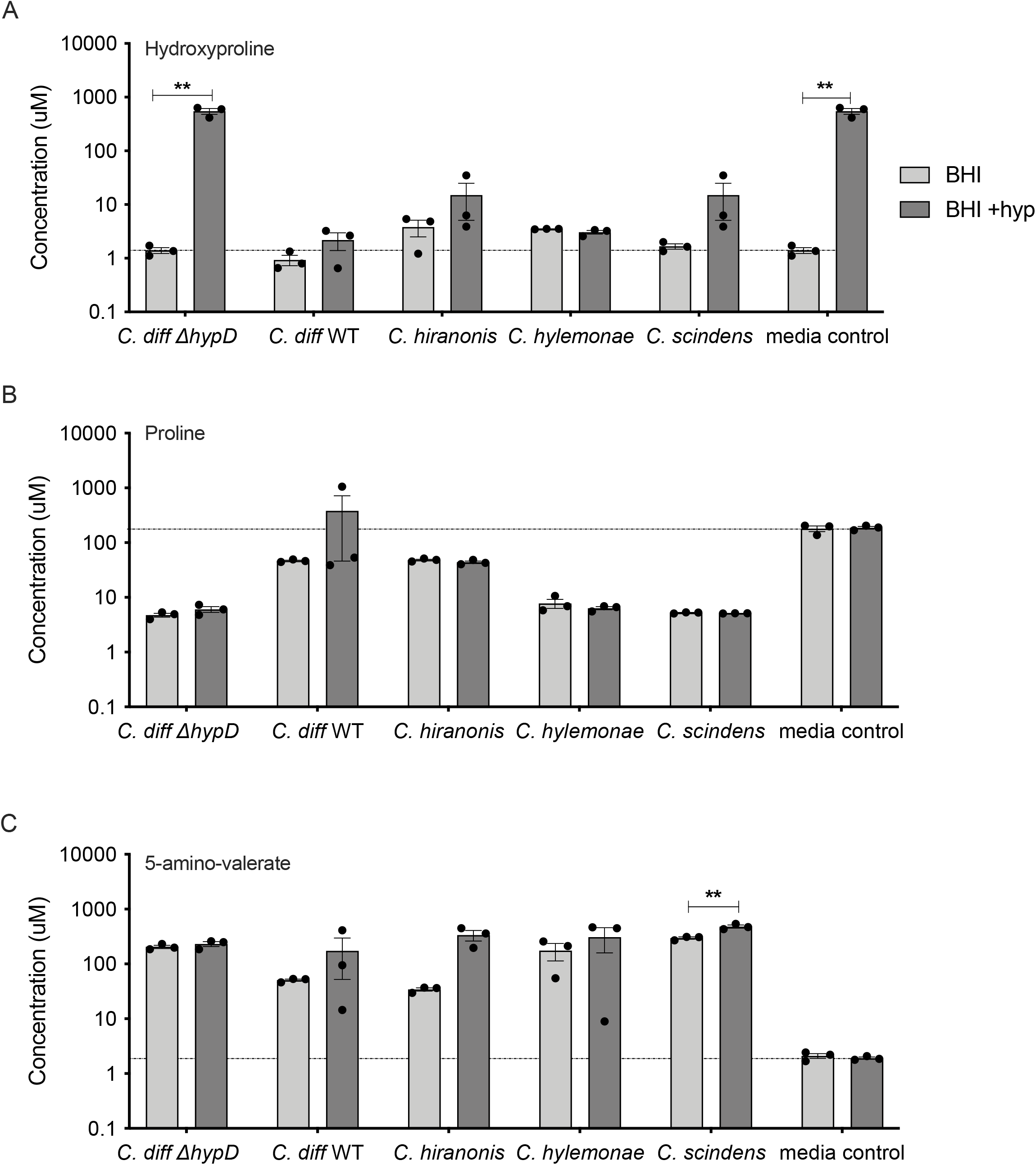
*C. difficile* WT, *C. hiranonis, C. hylemonae*, and *C. scindens* utilize hydroxyproline when it is supplemented into a rich growth medium. Concentration of (**A**) hydroxyproline, (**B**) proline, and (**C**) 5-amino-valerate in BHI and in BHI +600 mg/L hydroxyproline. Supernatants were taken after 24 hours of growth by WT, *hypD, C. hiranonis, C. hylemonae*, or *C. scindens.* BHI alone (dotted line) and BHI +600mg/L of hydroxyproline were used as controls. Statistical significance was determined using Student’s T Test with Welch’s Correction to account for multiple comparisons (*, P<0.05; **, P<0.01; ***, P<0.001; ****, P<0.0001).

### The genomic position of *p5cr* in relation to *hypD* and the transcriptional response to hydroxyproline varies between *C. difficile* and commensal *Clostridia*

When *hypD* (CD630_32820) and *p5cr* (CD630_32810) were aligned across strains using *C. difficile* as the reference strain, it was found that only *C. difficile* and *C. hiranonis* had the *p5cr* gene next to the *hypD* gene (Figure 5A). In *C. hylemonae* and *C. scindens*, the *p5cr* gene was not adjacent to the *hypD* gene. In addition, *hypD from C. hiranonis* showed the greatest amino acid identity (84%) to the *C. difficile* HypD protein while the other two commensals only showed 55% (Figure 5A). All commensals showed a 69% AA identity to the P5CR protein encoded by *C. difficile.* To determine if this would affect the transcriptional response of *hypD* and *p5cr* to hydroxyproline, *C. difficile, C. hiranonis, C. hylemonae* and *C. scindens* were each grown in BHI or BHI + hyp overnight and the relative copy number of *hypD* and *p5cr* transcripts were analyzed using qRT-PCR (Figure 5B–E).

**Figure 5:**
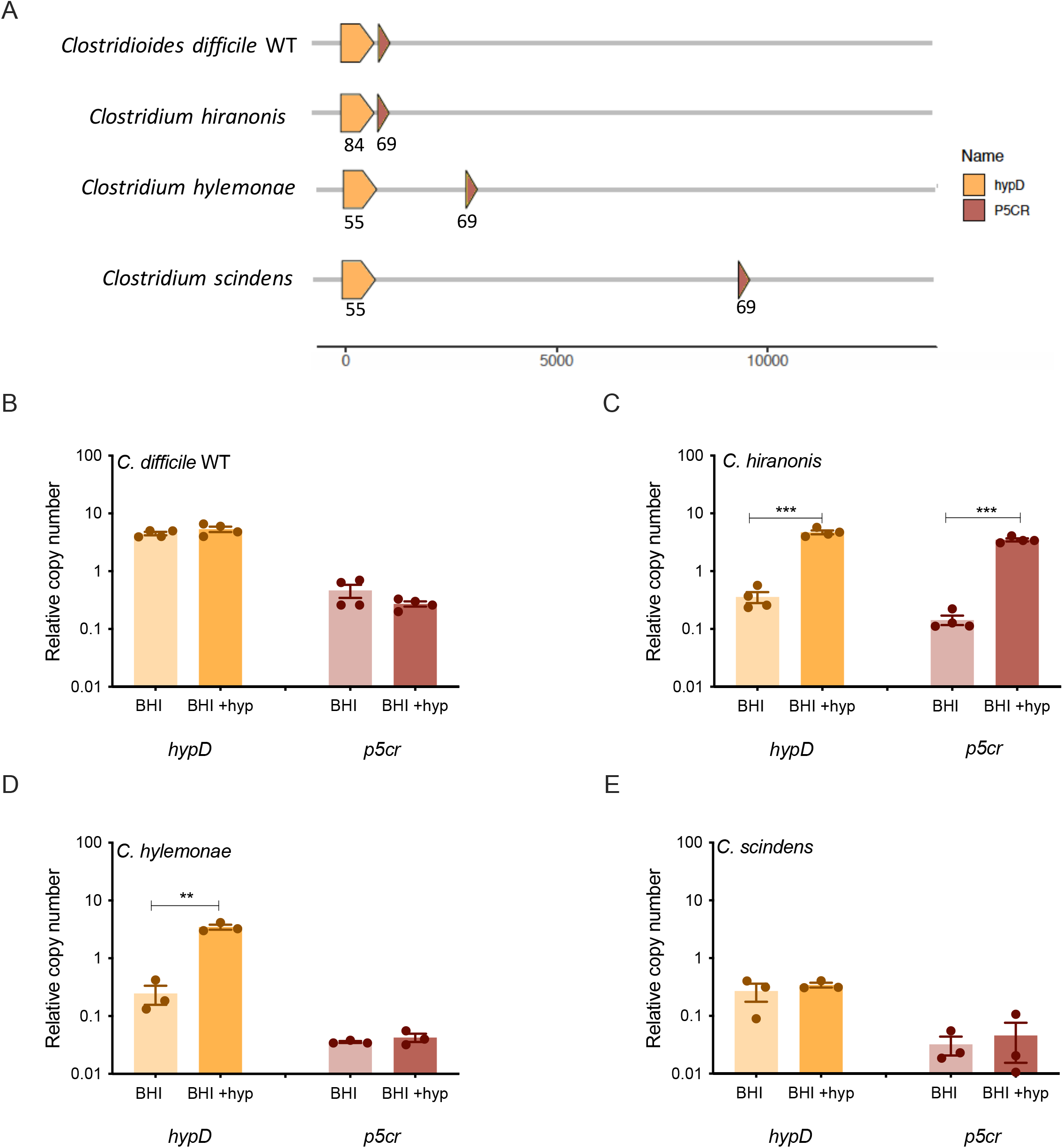
Expression of *hypD* and *p5cr* differs between *C. difficile* and selected commensal *Clostridia* in rich media supplemented with hydroxyproline. **A.** Alignment of *hypD* and *p5cr* across *C. difficile* and selected commensal *Clostridia* strains. Each protein sequence was compared against its counterpart in the reference strain *C. difficile* 630Δ*erm*, generating the amino acid percent identity labeled within each gene. **B-E**. Expression of *hypD* and *p5cr* in BHI media and BHI media with 600 mg/L hydroxyproline added of *C. difficile* (**B**)*, C. hiranonis* (**C**), *C. hylemonae* (**D**) and *C. scindens* (**E**). Experiments were run in triplicate and three biological replicates were performed. The expression in medium supplemented with hydroxyproline was compared to expression in medium without additional hydroxyproline. Statistical significance was determined using Student’s T Test (*, P<0.05; **, P<0.01; ***, P<0.001; ****, P<0.0001).

All four strains tested had different transcriptional responses to hydroxyproline supplementation of rich media (Figure 5B–E). *C. hiranonis* had significantly increased expression of *hypD* and *p5cr* in the presence of hydroxyproline, which was expected given that the two genes are possibly operonic in that strain (p < 0.001, Student’s T test). *C. hylemonae* had significantly increased expression of *hypD*, but not of *p5cr* (p < 0.01, Student’s T test). Neither *C. difficile* nor *C. scindens* showed significantly altered expression for either gene, but the overall relative copy number for *C. difficile* was approximately ten-fold higher than the relative copy number for *C. scindens* (Figure 5B, 5E).

### *C. difficile* and commensal *Clostridia* each have different transcriptomic responses to the presence of hydroxyproline

*C. difficile*, *C. hiranonis, C. hylemonae* and *C. scindens* were all grown in BHI or BHI supplemented with 600mg/L of hydroxyproline (BHI + hyp) media. At mid-log growth (OD_600_ 0.3-0.5), RNA was extracted, and the transcriptomic response was analyzed using RNAseq. For *C. difficile*, many of the genes that were upregulated upon exposure to hydroxyproline are involved in proline metabolism, including the copy of *proC* that is adjacent to *hypD*. In particular, many of the genes in the *prd* operon, which encodes enzymes for the reduction of proline in Stickland fermentation and has previously been shown to be upregulated in the presence of proline, were upregulated in *C. difficile* (Figure 6A, Supplemental Data File 4)[26]. Genes involved in regenerating NAD+ via the reduction of succinate and its conversion to butyrate were decreased in expression in the presence of hydroxyproline, consistent with the role of proline reductase as a preferred mechanism of reducing equivalent regeneration. In *C. hiranonis*, most of the differentially expressed genes were downregulated, including amino acid and branched chain amino acid biosynthetic genes, as well as carbohydrate utilization genes. The putative ferrous iron importer gene*feoB2* was increased in *C. hiranonis* in the presence of hydroxyproline, as well as gene encoding a putative NADP-dependent *α*-hydroxysteroid dehydrogenase, although the overall expression of the latter was quite low (Fig. 6B). Similarly, *C. hylemonae* had several transcripts that significantly decreased with supplementation of hydroxyproline, including those encoding the glycine reductase (Fig. 6C). Conversely, the genes encoding the glycine cleavage system were increased in *C. hiranonis* in the presence of hydroxyproline. *C. scindens* had the largest number of differentially expressed genes between the two media conditions (Figure 6D). A number of genes from the *prd* operon were upregulated in response to hydroxyproline, as were a number of genes encoding subunits of an electron transport complex (*rnfABCDGE*) (Figure 6D). Several genes from the *bai* (bile acid inducible) operon were significantly decreased, although their expression levels in BHI alone were quite low. The expression of genes in the *bai* operon was decreased in *C. scindens* and *C. hylemonae*, in the presence of hydroxyproline but in *C. hiranonis*, the expression of *bai* operon genes was increased in the presence of hydroxyproline (Supplemental Figure 3). Overall, the variable transcriptional responses to the presence of hydroxyproline observed between *C. difficile* and the three commensal *Clostridia* revealed changes in non-hydroxyproline associated metabolic pathways, including those for fermentation of other Stickland substrates.

**Figure 6:**
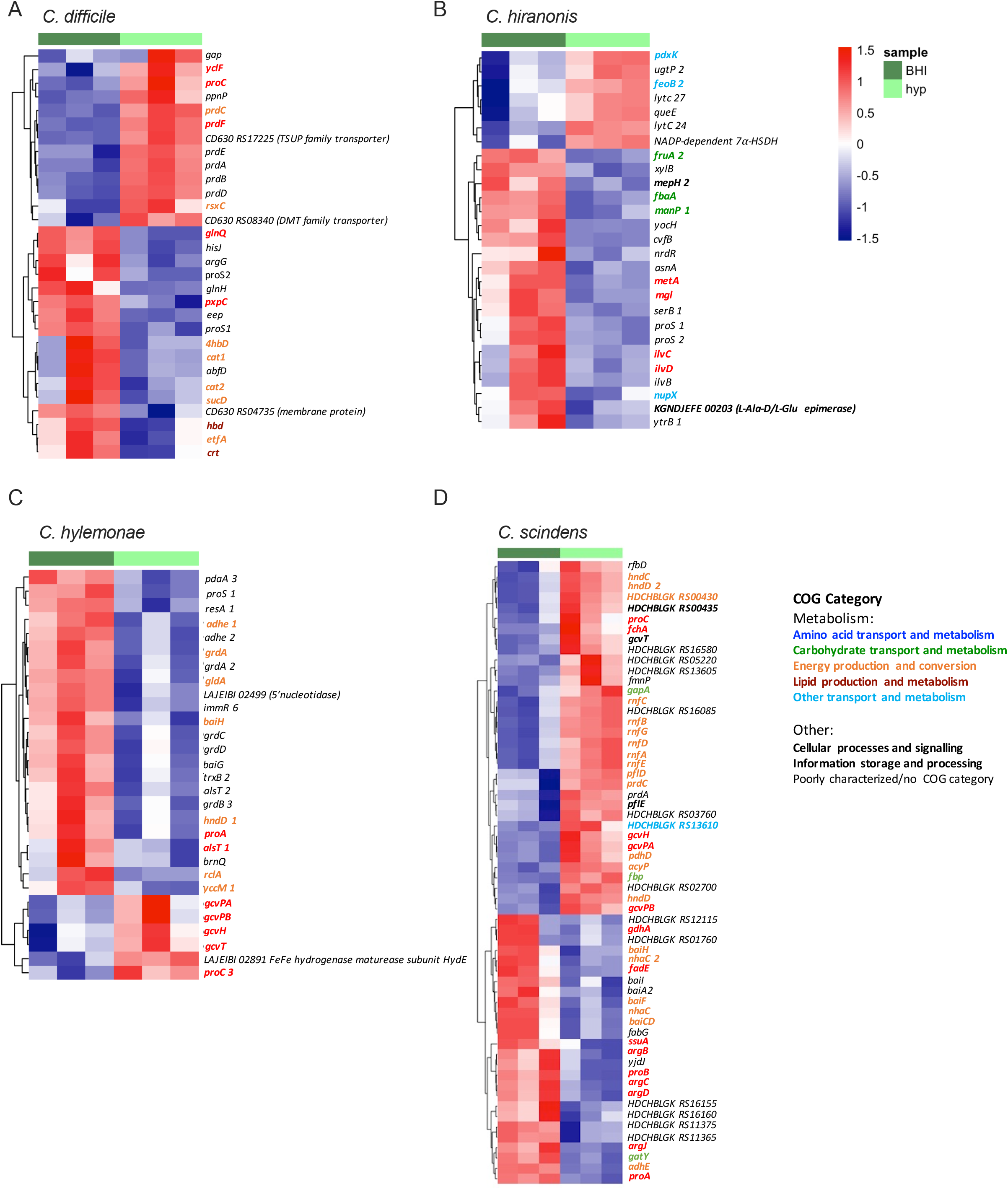
Transcriptomic differences in response to hydroxyproline vary between *C. difficile* and commensal *Clostridia*. Heatmap of genes that had significantly differential expression in **(A)** *C. difficile* WT, **(B)** *C. hiranonis*, **(C)** *C. hylemonae* and **(D)** *C. scindens* between Analysis was run using Geneious and DESeq2. All genes considered differentially expressed had an adjusted p value of <0.05 and ±1 log fold change.

## Discussion

Understanding which nutrients are required for *C. difficile* to persist and cause disease in the host is important to developing targeted therapeutics against CDI. In this study, we employed bacterial genetics to examine how the utilization of hydroxyproline by *C. difficile* affects CDI and the microbiome in a mouse model of infection. To facilitate the interpretation of the *in vivo* data, the Δ*hypD* and Δ*p5cr* mutants and their complements were first grown in a minimal medium with hydroxyproline substituted for proline (CDMM –pro). While there was a growth defect in the Δ*hypD* mutant, as expected, no growth defect was observed for the Δ*p5cr* mutant (Figure 1B). This is potentially due to functional redundancy within the *C. difficile630Δerm* genome where a second homolog of P5CR is present (encoded by *CD630_14950*), which did not significantly change expression in the presence of hydroxyproline. Also of interest is that all strains other than the Δ*hypD* mutant showed significantly increased growth when hydroxyproline was present in the media as opposed to proline. This is likely due to the repression of alternative NAD^+^ regeneration pathways that would result in butyrate production, which would impede growth of *C. difficile*, but further experiments are required to test this hypothesis[25,29,40].

Since Δ*hypD* growth was impaired *in vitro* when hydroxyproline was the only proline source, we reasoned that *hypD* may be important for colonization and disease progression in a mouse model of CDI. While the fecal burden of *C. difficile* was similar between the strains, the mice challenged with the WT strain showed more weight loss on days 3, 5 and 7 after challenge and toxin activity was higher on day 3 post challenge relative to mice colonized with Δ*hypD* (Figure 2D–E). There was no significant difference in toxin activity 7 days post challenge, indicating that this effect is the strongest earlier during disease. While the effect is subtle, the differences in weight and toxin activity suggest that *C. difficile* relies on hydroxyproline for maximal fitness *in vivo*, highlighting the importance of a host-derived amino acid that is likely made available via toxin-induced expression of host matrix metalloproteinases[14,41,42]. *C. difficile* 630 was chosen for this experiment due to the genetic tools available for this strain, but it causes less severe disease in a mouse model than strains R20291 or VPI 10463 [43–45]. It is possible that a stronger difference between the mutant and the wild type strain would be observed in these strain backgrounds, especially given their increased expression of *hypD* in the presence of hydroxyproline in a defined medium when compared to *C. difficile* 630 (Supplemental Figure 4).

Each of the bacteria tested had a different transcriptional response to hydroxyproline, both in terms of RNAseq and when *hypD* and *p5cr* were tested individually using qRT-PCR (Figures 5–6). Of particular interest was the fact that neither *C. difficile* nor *C. scindens* showed upregulation of *hypD* or *p5cr* when hydroxyproline was supplemented to the media but when the levels of amino acids were quantified using LC/MS, both organisms metabolized the majority of hydroxyproline present (Figure 4A). For *C. scindens*, this may mean that *hypD* and/or *p5cr* are always transcriptionally active or that the bacterium has another way to utilize hydroxyproline that doesn’t require either gene. For *C. difficile* 630*Δerm*, it is more likely that *hypD* is always transcriptionally active, as *C. difficile* Δ*hypD* did not utilize the excess hydroxyproline added to the media, in addition to the growth defect previously observed (Figure 1B and 4A). The lack of differential expression in *C. difficile* 630 is particularly interesting, as when the *C. difficile* strains 630, R20291 and VPI 10463 were tested in a minimal medium, 630 was the only one where *hypD* was not strongly upregulated in the presence of hydroxyproline, indicating that there are regulatory differences between strains (Supplemental Figure 4). Unfortunately, one of the limitations of the *in vitro* work in this study was the requirement to use a rich and undefined medium, that contains a basal level of hydroxyproline, as *C. hiranonis* and *C. hylemonae* do not grow well in defined media[16,46].

The overall transcriptional response of *C. difficile* 630Δ*erm* and the commensal *Clostridia* to hydroxyproline indicated *in vitro* that while there were some similarities, each organism had a relatively unique response. In *C. scindens*, over 60 transcripts were significantly altered in response to hydroxyproline, with 38 of those genes being in a metabolic COG category. Of particular interest is that *baiA2, baiCD, baiF and baiH* were all downregulated in response to hydroxyproline, indicating that even without cholate in the media, the activation of the *bai* operon can vary depending on the nutritional content of the media. While none of the changes in *bai* transcripts in *C. hylemonae* were statistically significant, several *bai* operon genes, including *baiG* and *baiE*, were downregulated in response to hydroxyproline (Supplemental Figure 3). This is particularly interesting given the previous finding that *C. hylemonae* shows upregulation of the *bai* operon when exposed to cholate in a defined medium, but not when exposed to cholate in BHI [16,46]. Despite recent work that suggests bile acids do not play an essential role in protection against CDI, this study provides further evidence that there is a relationship between nutrient availability and secondary bile acid production in commensal *Clostridia* [47]. Further work combining bile acids and hydroxyproline, and other amino acids important for Stickland fermentation, are needed to fully dissect transcriptional networks in these organisms and define their individual and combinatorial roles in colonization resistance against *C. difficile*. This supports the finding that each of these commensal *Clostridia* have differing metabolic responses to hydroxyproline, and that further elucidation of their nutrient utilization *in vivo* will be fruitful for identifying possible nutritional overlaps with *C. difficile.* This approach may allow for the development of rationally designed cocktails of commensal microbiota that can compete against *C. difficile* for one or more nutrient sources in an infected host.

## Acknowledgements

We would like to thank Jason Ridlon for providing the commensal *Clostridia* strains. YYH, DJK and EPB were funded by Faculty Scholar Award and the Packard Fellowship for Science and Engineering from the David and Lucile Packard Foundation (2013-39267). CMT was funded by the National Institute of General Medical Sciences of the National Institutes of Health under award number R35GM119438. This work was performed in part by the Molecular Education, Technology and Research Innovation Center (METRIC) at NC State University, which is supported by the State of North Carolina.

## Data availability

Raw sequences have been deposited in the Sequence Read Archive (SRA) with SRA submission number SUB10603241 for 16S data and SUB10603493 for RNAseq data. They can both be found under BioProject ID PRJNA776739. Source data are provided within each Supplementary Data Files. Other data and biological materials are available from the corresponding author upon reasonable requests.

---

**Supplemental Figure 1: Vegetative bacterial load in cecal content is higher in Δ*hypD* mutant than in WT on Day 7.** *C. difficile* vegetative cell (**A**) or spore (**B**) CFUs in cecal content on day 7 post challenge. Statistical significance was determined using Mann-Whitney (*, p<0.05; **, p<0.01; ***, p<0.001; ****, p<0.0001).

**Supplemental Figure 2: The Alpha and Beta diversity differ between groups on day 7. A.** Alpha diversity calculated using Inverse simpson at the family level for cef, *hypD* and WT fecal microbiome for days 0, 2, 4, and 6 post challenge and the cecal microbiome for day 7 post challenge. **B.** Beta diversity calculated using NMDS for cef, *hypD* and WT fecal microbiome for days 0, 2, 4, and 6 post challenge and the cecal microbiome for day 7 post challenge. **C.** Beta diversity calculated using NMDS for *hypD* and WT fecal microbiome for days 0, 2, 4, and 6 post challenge and the cecal microbiome for day 7 post challenge. Statistical significance was determined using Kruskal-Wallis for Alpha diversity and Permanova Adonis for Beta diversity (*, p<0.05; **, p<0.01; ***, p<0.001; ****, p<0.0001).

**Supplemental Figure 3. The transcriptional response of *hypD* to hydroxyproline differs between *C. difficile* strains.** Expression of *hypD* in CDMM and CDMM –pro +hyp of *C. difficile* 630, R20291 and VPI 10463. Experiments were run in triplicate and two biological replicates were performed.

**Supplemental Figure 4: Transcriptional response of *bai* operon to hydroxyproline differs between *C. hiranonis* and other commensal *Clostridia.*** Heatmap of *baiA1* and genes within the *bai* operon in BHI and BHI + hyp in **(A)** *C. hiranonis*,**(B)** *C. hylemonae* and **(C)** *C. scindens.*

## References

1. Lessa, F.C.; Mu, Y.; Bamberg, W.M.; Beldavs, Z.G.; Dumyati, G.K.; Dunn, J.R.; Farley, M.M.; Holzbauer, S.M.; Meek, J.I.; Phipps, E.C., et al. Burden of *Clostridium difficile* infection in the United States. N Engl J Med 2015, 372, 825–834, doi:10.1056/NEJMoa1408913.

2. Magill, S.S.; O’Leary, E.; Janelle, S.J.; Thompson, D.L.; Dumyati, G.; Nadle, J.; Wilson, L.E.; Kainer, M.A.; Lynfield, R.; Greissman, S., et al. Changes in Prevalence of Health Care-Associated Infections in U.S. Hospitals. N Engl J Med 2018, 379, 1732–1744, doi:10.1056/NEJMoa1801550.

3. Kelly, C.P. Current strategies for management of initial Clostridium difficile infection. J Hosp Med 2012, 7 Suppl 3, S5–10, doi:10.1002/jhm.1909.

4. Cornely, O.A.; Miller, M.A.; Louie, T.J.; Crook, D.W.; Gorbach, S.L. Treatment of first recurrence of Clostridium difficile infection: fidaxomicin versus vancomycin. Clin Infect Dis 2012, 55 Suppl 2, S154–161, doi:10.1093/cid/cis462.

5. Fekety, R.; McFarland, L.V.; Surawicz, C.M.; Greenberg, R.N.; Elmer, G.W.; Mulligan, M.E. Recurrent Clostridium difficile diarrhea: characteristics of and risk factors for patients enrolled in a prospective, randomized, double-blinded trial. Clin Infect Dis 1997, 24, 324–333, doi:10.1093/clinids/24.3.324.

6. Owens, R.C., Jr.; Donskey, C.J.; Gaynes, R.P.; Loo, V.G.; Muto, C.A. Antimicrobial-associated risk factors for Clostridium difficile infection. Clin Infect Dis 2008, 46 Suppl 1, S19–31, doi:10.1086/521859.

7. Seekatz, A.M.; Young, V.B. Clostridium difficile and the microbiota. J Clin Invest 2014, 124, 4182–4189, doi:10.1172/JCI72336.

8. Theriot, C.M.; Koenigsknecht, M.J.; Carlson, P.E., Jr.; Hatton, G.E.; Nelson, A.M.; Li, B.; Huffnagle, G.B.; J, Z.L.; Young, V.B. Antibiotic-induced shifts in the mouse gut microbiome and metabolome increase susceptibility to Clostridium difficile infection. Nat Commun 2014, 5, 3114, doi:10.1038/ncomms4114.

9. Ducarmon, Q.R.; Zwittink, R.D.; Hornung, B.V.H.; van Schaik, W.; Young, V.B.; Kuijper, E.J. Gut Microbiota and Colonization Resistance against Bacterial Enteric Infection. Microbiology and Molecular Biology Reviews 2019, 83, e00007–00019, doi:10.1128/mmbr.00007-19.

10. Crobach, M.J.T.; Vernon, J.J.; Loo, V.G.; Kong, L.Y.; Pechine, S.; Wilcox, M.H.; Kuijper, E.J. Understanding *Clostridium difficile* Colonization. Clin Microbiol Rev 2018, 31, doi:10.1128/CMR.00021-17.

11. Buffie, C.G.; Pamer, E.G. Microbiota-mediated colonization resistance against intestinal pathogens. Nature Reviews Immunology 2013, 13, 790–801.

12. Guo, C.J.; Allen, B.M.; Hiam, K.J.; Dodd, D.; Van Treuren, W.; Higginbottom, S.; Nagashima, K.; Fischer, C.R.; Sonnenburg, J.L.; Spitzer, M.H., et al. Depletion of microbiome-derived molecules in the host using Clostridium genetics. Science 2019, 366, doi:10.1126/science.aav1282.

13. Battaglioli, E.J.; Hale, V.L.; Chen, J.; Jeraldo, P.; Ruiz-Mojica, C.; Schmidt, B.A.; Rekdal, V.M.; Till, L.M.; Huq, L.; Smits, S.A., et al. Clostridioides difficile uses amino acids associated with gut microbial dysbiosis in a subset of patients with diarrhea. Sci Transl Med 2018, 10, doi:10.1126/scitranslmed.aam7019.

14. Fletcher, J.R.; Pike, C.M.; Parsons, R.J.; Rivera, A.J.; Foley, M.H.; McLaren, M.R.; Montgomery, S.A.; Theriot, C.M. Clostridioides difficile exploits toxin-mediated inflammation to alter the host nutritional landscape and exclude competitors from the gut microbiota. Nature Communications 2021, 12, 1–14.

15. Fletcher, J.R.; Erwin, S.; Lanzas, C.; Theriot, C.M. Shifts in the Gut Metabolome and Clostridium difficile Transcriptome throughout Colonization and Infection in a Mouse Model. mSphere 2018, 3, doi:10.1128/mSphere.00089-18.

16. Reed, A.D.; Nethery, M.A.; Stewart, A.; Barrangou, R.; Theriot, C.M. Strain-dependent inhibition of Clostridioides difficile by commensal Clostridia encoding the bile acid inducible (bai) operon. J Bacteriol 2020, 10.1128/JB.00039-20, doi:10.1128/JB.00039-20.

17. Winston, J.A.; Rivera, A.J.; Cai, J.; Thanissery, R.; Montgomery, S.A.; Patterson, A.D.; Theriot, C.M. Ursodeoxycholic Acid (UDCA) Mitigates the Host Inflammatory Response during Clostridioides difficile Infection by Altering Gut Bile Acids. Infect Immun 2020, 88, doi:10.1128/IAI.00045-20.

18. Ridlon, J.M.; Harris, S.C.; Bhowmik, S.; Kang, D.J.; Hylemon, P.B. Consequences of bile salt biotransformations by intestinal bacteria. Gut Microbes 2016, 7, 22–39, doi:10.1080/19490976.2015.1127483.

19. Buffie, C.G.; Bucci, V.; Stein, R.R.; McKenney, P.T.; Ling, L.; Gobourne, A.; No, D.; Liu, H.; Kinnebrew, M.; Viale, A., et al. Precision microbiome reconstitution restores bile acid mediated resistance to Clostridium difficile. Nature 2015, 517, 205–208, doi:10.1038/nature13828.

20. Sorg, J.A.; Sonenshein, A.L. Inhibiting the initiation of Clostridium difficile spore germination using analogs of chenodeoxycholic acid, a bile acid. J Bacteriol 2010, 192, 4983–4990, doi:10.1128/JB.00610-10.

21. Wilson, K.H.; Sheagren, J.N. Antagonism of toxigenic Clostridium difficile by nontoxigenic C. difficile. J Infect Dis 1983, 147, 733–736, doi:10.1093/infdis/147.4.733.

22. Gerding, D.N.; Meyer, T.; Lee, C.; Cohen, S.H.; Murthy, U.K.; Poirier, A.; Van Schooneveld, T.C.; Pardi, D.S.; Ramos, A.; Barron, M.A., et al. Administration of spores of nontoxigenic Clostridium difficile strain M3 for prevention of recurrent C. difficile infection: a randomized clinical trial. JAMA 2015, 313, 1719–1727, doi:10.1001/jama.2015.3725.

23. Jenior, M.L.; Leslie, J.L.; Powers, D.A.; Garrett, E.M.; Walker, K.A.; Dickenson, M.E.; Petri, W.A.; Tamayo, R.; Papin, J.A. Novel drivers of virulence in Clostridioides difficile identified via context-specific metabolic network analysis. bioRxiv 2021, 2020.2011.2009.373480.

24. Jenior, M.L.; Leslie, J.L.; Young, V.B.; Schloss, P.D. Clostridium difficile colonizes alternative nutrient niches during infection across distinct murine gut microbiomes. Msystems 2017, 2.

25. Bouillaut, L.; Self, W.T.; Sonenshein, A.L. Proline-dependent regulation of Clostridium difficile Stickland metabolism. J Bacteriol 2013, 195, 844–854, doi:10.1128/JB.01492-12.

26. Karasawa, T.; Ikoma, S.; Yamakawa, K.; Nakamura, S. A defined growth medium for Clostridium difficile. Microbiology 1995, 141 (Pt 2), 371–375, doi:10.1099/13500872-141-2-371.

27. Gencic, S.; Grahame, D.A. Diverse energy-conserving pathways in Clostridium difficile: Growth in the absence of amino acid Stickland acceptors and the role of the Wood-Ljungdahl pathway. Journal of bacteriology 2020, 202, e00233–00220.

28. Gorres, K.L.; Raines, R.T. Prolyl 4-hydroxylase. Critical reviews in biochemistry and molecular biology 2010, 45, 106–124.

29. Huang, Y.Y.; Martinez-Del Campo, A.; Balskus, E.P. Anaerobic 4-hydroxyproline utilization: Discovery of a new glycyl radical enzyme in the human gut microbiome uncovers a widespread microbial metabolic activity. Gut Microbes 2018, 9, 437–451, doi:10.1080/19490976.2018.1435244.

30. Levin, B.J.; Huang, Y.Y.; Peck, S.C.; Wei, Y.; Martinez-Del Campo, A.; Marks, J.A.; Franzosa, E.A.; Huttenhower, C.; Balskus, E.P. A prominent glycyl radical enzyme in human gut microbiomes metabolizes trans-4-hydroxy-l-proline. Science 2017, 355, doi:10.1126/science.aai8386.

31. Lopez, C.A.; McNeely, T.P.; Nurmakova, K.; Beavers, W.N.; Skaar, E.P. Clostridioides difficile proline fermentation in response to commensal clostridia. Anaerobe 2020, 63, 102210, doi:10.1016/j.anaerobe.2020.102210.

32. Thanissery, R.; Winston, J.A.; Theriot, C.M. Inhibition of spore germination, growth, and toxin activity of clinically relevant C. difficile strains by gut microbiota derived secondary bile acids. Anaerobe 2017.

33. Thanissery, R.; Zeng, D.; Doyle, R.G.; Theriot, C.M. A Small Molecule-Screening Pipeline to Evaluate the Therapeutic Potential of 2-Aminoimidazole Molecules Against Clostridium difficile. Front Microbiol 2018, 9, 1206, doi:10.3389/fmicb.2018.01206.

34. Gibson, D.G.; Young, L.; Chuang, R.Y.; Venter, J.C.; Hutchison, C.A., 3rd; Smith, H.O. Enzymatic assembly of DNA molecules up to several hundred kilobases. Nat Methods 2009, 6, 343–345, doi:10.1038/nmeth.1318.

35. Fimlaid, K.A.; Jensen, O.; Donnelly, M.L.; Francis, M.B.; Sorg, J.A.; Shen, A. Identification of a Novel Lipoprotein Regulator of Clostridium difficile Spore Germination. PLoS Pathogens 2015, 10.1371/journal.ppat.1005239, doi:10.1371/journal.ppat.1005239.

36. Putnam, E.E.; Nock, A.M.; Lawley, T.D.; Shen, A. SpoIVA and SipL are Clostridium difficile spore morphogenetic proteins. J Bacteriol 2013, 195, 1214–1225, doi:10.1128/JB.02181-12.

37. Gloor, G. ALDEx2: ANOVA-Like Differential Expression tool for compositional data. ALDEX manual modular 2015, 20, 1–11.

38. Gómez-Rubio, V. ggplot2-elegant graphics for data analysis. Journal of Statistical Software 2017, 77, 1–3.

39. McMurdie, P.J.; Holmes, S. phyloseq: an R package for reproducible interactive analysis and graphics of microbiome census data. PloS one 2013, 8, e61217.

40. Neumann-Schaal, M.; Jahn, D.; Schmidt-Hohagen, K. Metabolism the Difficile Way: The Key to the Success of the Pathogen Clostridioides difficile. Front Microbiol 2019, 10, 219, doi:10.3389/fmicb.2019.00219.

41. Hensbergen, P.J.; Klychnikov, O.I.; Bakker, D.; Dragan, I.; Kelly, M.L.; Minton, N.P.; Corver, J.; Kuijper, E.J.; Drijfhout, J.W.; van Leeuwen, H.C. *Clostridium difficile* secreted Pro-Pro endopeptidase PPEP-1 (ZMP1/CD2830) modulates adhesion through cleavage of the collagen binding protein CD2831. FEBS Lett 2015, 589, 3952–3958, doi:10.1016/j.febslet.2015.10.027.

42. Udenfriend, S. Formation of hydroxyproline in collagen. Science 1966, 152, 1335–1340.

43. Winston, J.A.; Thanissery, R.; Montgomery, S.A.; Theriot, C.M. Cefoperazone-treated Mouse Model of Clinically-relevant Clostridium difficile Strain R20291. J Vis Exp 2016, 10.3791/54850, doi:10.3791/54850.

44. Koenigsknecht, M.J.; Theriot, C.M.; Bergin, I.L.; Schumacher, C.A.; Schloss, P.D.; Young, V.B. Dynamics and establishment of Clostridium difficile infection in the murine gastrointestinal tract. Infect Immun 2015, 83, 934–941, doi:10.1128/IAI.02768-14.

45. Theriot, C.M.; Koumpouras, C.C.; Carlson, P.E.; Bergin, I.I.; Aronoff, D.M.; Young, V.B. Cefoperazone-treated mice as an experimental platform to assess differential virulence of Clostridium difficile strains. Gut microbes 2011, 2, 326–334.

46. Ridlon, J.M.; Devendran, S.; Alves, J.M.; Doden, H.; Wolf, P.G.; Pereira, G.V.; Ly, L.; Volland, A.; Takei, H.; Nittono, H., et al. The ‘in vivo lifestyle’ of bile acid 7alpha-dehydroxylating bacteria: comparative genomics, metatranscriptomic, and bile acid metabolomics analysis of a defined microbial community in gnotobiotic mice. Gut Microbes 2019, 10.1080/19490976.2019.1618173, 1–24, doi:10.1080/19490976.2019.1618173.

47. Aguirre, A.M.; Yalcinkaya, N.; Wu, Q.; Swennes, A.; Tessier, M.E.; Roberts, P.; Miyajima, F.; Savidge, T.; and Sorg, J.A. Bile acid-independent protection against Clostridioides difficile infection. PLoS Pathogens 17(10): e1010015.

